# The Precision Medicine Knowledge Base: an online application for collaborative editing, maintenance and sharing of structured clinical-grade cancer mutations interpretations

**DOI:** 10.1101/059824

**Authors:** Linda Huang, Helen Fernandes, Hamid Zia, Peyman Tavassoli, Hanna Rennert, David Pisapia, Marcin Imielinski, Andrea Sboner, Mark A. Rubin, Michael Kluk, Olivier Elemento

**Affiliations:** Institute for Precision Medicine Weill Cornell Medicine 1300 York Avenue New York, NY, 10021; Institute for Computational Biomedicine Weill Cornell Medicine 1300 York Avenue New York, NY, 10021; Department of Pathology and Laboratory Medicine Weill Cornell Medicine 1300 York Avenue New York, NY, 10021

**Keywords:** precision medicine, database, cancer genomics, clinical reporting, pathology, application-programming interface

## Abstract

**Objective:** This paper describes the Precision Medicine Knowledge Base (PMKB; https://pmkb.weill.cornell.edu), an interactive online application for collaborative editing, maintenance and sharing of structured clinical-grade cancer mutations interpretations.

**Materials and Methods:** PMKB was built using the Ruby on Rails Web application framework. Leveraging existing standards such as Human Genome Variation Society (HGVS) variant description format, we implemented a data model that links variants to tumor-specific and tissue-specific interpretations. Key features of PMKB include support for all major variant types, standardized authentication, distinct user roles including high-level approvers, detailed activity history. A REpresentational State Transfer (REST) application-programming interface (API) was implemented to query the PMKB programmatically.

**Results:** At the time of writing, PMKB contains 457 variant descriptions with 281 clinical-grade interpretations. The EGFR, BRAF, KRAS, and KIT genes are associated with the largest numbers of interpretable variants. The PMKB’s interpretations have been used in over 1,500 AmpliSeq tests and 750 whole exome sequencing tests. The interpretations are accessed either directly via the Web interface or programmatically via the existing API.

**Discussion:** An accurate and up-to-date knowledge base of genomic alterations of clinical significance is critical to the success of precision medicine programs. The open-access, programmatically accessible PMKB represents an important attempt at creating such a resource in the field of oncology.

**Conclusion:** The PMKB was designed to help collect and maintain clinical-grade mutation interpretations and facilitates reporting for clinical cancer genomic testing. The PMKB was also designed to enable the creation of clinical cancer genomics automated reporting pipelines via an API.

## BACKGROUND AND SIGNIFICANCE

A growing number of medical institutions have started genomic testing-driven precision medicine programs (1–3). A critical component of clinical genomic testing is the generation of accurate and informative reports containing clinical-grade interpretations of genomic alterations. Such reports must not only list which variants and mutations were found in a given clinical sample but also provide an interpretation of these variants in the context of available and relevant clinical information. In cancer, the clinical significance and interpretation of somatic mutations and germline variants often depends on the tumor context, i.e., tumor type and site. At Weill-Cornell Medicine’s (WCM) Institute for Precision Medicine (IPM), together with New York Presbyterian Hospital (NYPH), we routinely conduct genomic sequencing using both targeted testing of hotspot mutations (AmpliSeq™ Cancer Hotspot Panel v2, Life Technologies) and whole exome sequencing (3). These tests are approved by the New York State Department of Health (NYS DOH). After sequencing and analyzing the data using either commercial (Torrent Suite, Life Technologies) or custom (whole-exome sequencing) pipelines, the position and effect of mutations on coding sequences are determined using publicly available tools including VEP(4), snpEff(5) and Annovar (6). Molecular pathologists who sign out genomic testing reports need to interpret the clinical relevance of each annotated mutation, summarizing their findings into a molecular report. This is usually a tedious task that requires extensive literature curation. Some resources such as ClinVar have started cataloguing the clinical significance for variants relative to disease phenotypes (7). To facilitate the task of interpreting cancer mutations, several online resources have attempted to curate and catalog clinically relevant mutations. These include Washington University’s CiViC DB (https://civic.genome.wustl.edu), MD Anderson’s Personalized Cancer Therapy (https://pct.mdanderson.org), Vanderbilt-Ingram Cancer Center’s MyCancerGenome (https://www.mycancergenome.org), and several others. While helpful resources, in our experience few of these databases contain clinical grade interpretations actually applicable to clinical reporting. In some instances, mutation interpretations do not meet required levels of brevity and specificity. In some databases, mutations are not interpreted in the context of specific tumor types. In others, only point mutations and indels are catalogued, while common clinically relevant mutations such as gene fusions and copy number alterations/variations (CNAs/CNVs) are not included. Several clinically critical features may be missing from these databases for integration into routine workflows, such as whether a variant is a pertinent negative in a given tumor type (that is, a variant for which information regarding the accuracy of negative calls must be reported). Some databases may have limited ability to maintain the information content up-to-date, or may lack versioning. Finally, while some databases have APIs for integration into automated workflows, most do not. To overcome these collective limitations, we created the Precision Medicine Knowledge Base. The PMKB is currently restricted to variant interpretations for oncology. It was designed in close collaboration with pathologists to ensure accurate and standardized terminology and workflows compatible with clinical use. Importantly, all interpretations are either written and/or approved by board-certified molecular pathologists. The PMKB’s interpretations have been used in over 1,500 AmpliSeq tests and 750 whole exome sequencing tests and are accessed either directly via a Web interface or programmatically via an API.

## MATERIALS AND METHODS

### Design

PMKB began development in 2015, in order to aid pathologist signout of AmpliSeq 50-gene panel results and was later expanded to support a broad array of features for signing out whole-exome sequencing reports such as copy number variations, germline variants, pertinent negatives and many more. PMKB is built using the Ruby on Rails web application framework, chosen for its popularity as a platform for complex moderate-load web applications, wide variety of open-source library extensions, and ease of use. Ruby on Rails provides interfacing with a relational database by default, providing users with the advantage of a database system that promotes data uniqueness and atomicity. PMKB uses a straightforward database schema that structures aspects of an interpretation in a modular way (**Figure S1**). The design considerations of PMKB's design and data fields are twofold: (1) to provide data granularity in such a way that makes automatic retrieval of interpretations possible, and (2) to provide a convenient experience for pathologists, who will be classifying variants and writing interpretations as part of clinical genomics sign-out.

### Data Granularity - Interpretations

An interpretation in PMKB requires associations to three different kinds of elements as part of its identity: (1) gene-variant descriptions, (2) cancer and tumor type descriptions, and (3) tissue type descriptions. The interpretation object itself contains the textual interpretation, supported by relevant literature, and a numeric tier. The tier is a category indicating how clinically actionable an interpretation is. This type of association captures the level of specificity expected in a clinical-grade report’s interpretations. When adding an interpretation to the PMKB, one can associate as many of each element (i.e., cancer/tumor and tissue types) as applicable to the interpretation. For example, an interpretation for a mutation in BRAF may be relevant to several different tissue types. This multiple-associations structure avoids repetition of interpretation text, since the mutation information is primarily linked to the cancer and tumor types. Associations can be as broad or as specific as necessary - for example, an interpretation could be specified for a variant in “any tumor type” or “any tissue” for more general comments. Separating the tumor type and tumor site also allows for more flexibility when generating new interpretations.

### Data Granularity - Variant Descriptions

We developed a system for variant description that incorporates existing standards such as that of the Human Genome Variation Society (HGVS) (8) (**Figure 1**). At the highest level, a variant is described by the gene it is associated to and a variant category. Variant categories include small, localized mutations (SNVs, indels), copy number alterations, and gene fusions. Descriptions of small, localized mutations such as single nucleotide variants (SNVs) and indels can be broken down into two groups: (1) A specific mutation described using HGVS protein-change and DNA-change notation, or (2) a gene region-based description. Gene regions can be further divided into (1) specific codons, (2) specific exons, and (3) the entire gene (**Figure 1**). The variant type, e.g. deletion, insertion, missense, nonsense, etc. must be specified. Fusions are described as a pair consisting of the primary gene and the partner gene. Copy number alterations are described as either a gain or loss of copy number, pertaining to either a gene or a chromosome region, using arm and cytoband notation, e.g. 17p13.1. When entering variants, PMKB automatically retrieves specific gene region information from Ensembl, based on Ensembl’s canonical transcript for a gene and Ensembl's GRCh37-based API (9).

**Figure 1:**
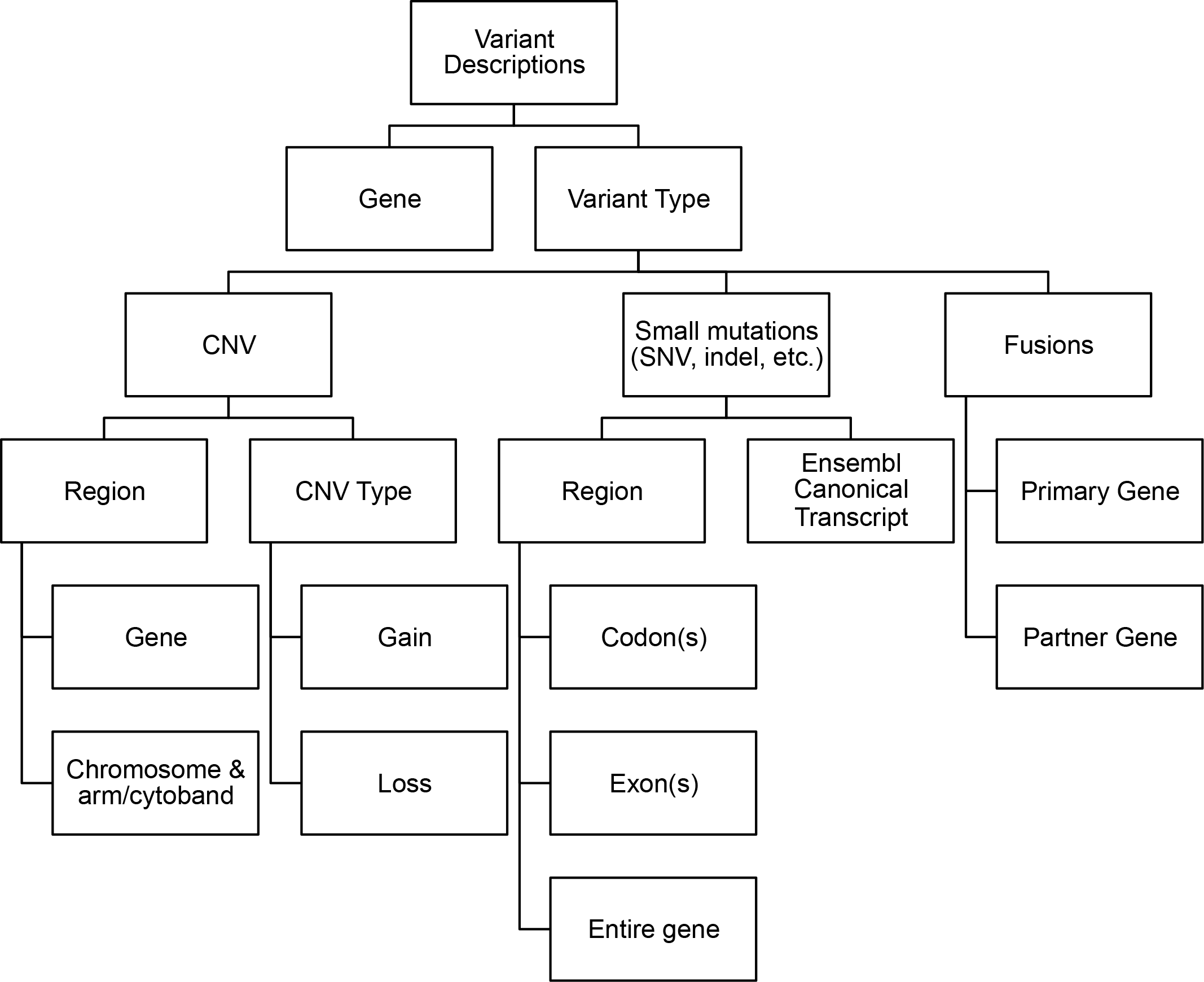
Diagram of the variant description data type.

Separating variant description into discrete fields facilitates the process of matching them against existing annotations. PMKB's REST API is set up to take a variant’s HGVS protein notation as input and match that variant against multiple levels of variant descriptions, then return all relevant interpretations. For example, in the case of a mutation, PMKB’s REST API would take KRAS p.G12A as input (**Figure 2**) and match it against all KRAS mutations in its database. This API query may return interpretations for KRAS p.G12A, KRAS codon 12 missense, KRAS exon 2 missense, KRAS exon 2 any mutation, and KRAS any mutation (**Figure 2**). These matches are ranked in order of specificity, based on the width of the sequence they fall in and whether the variant type is a match or ‘any’. Extra fields in the search query can make PMKB return only those interpretations that are linked to a specific tumor type and tissue type, if desired.

**Figure 2:**
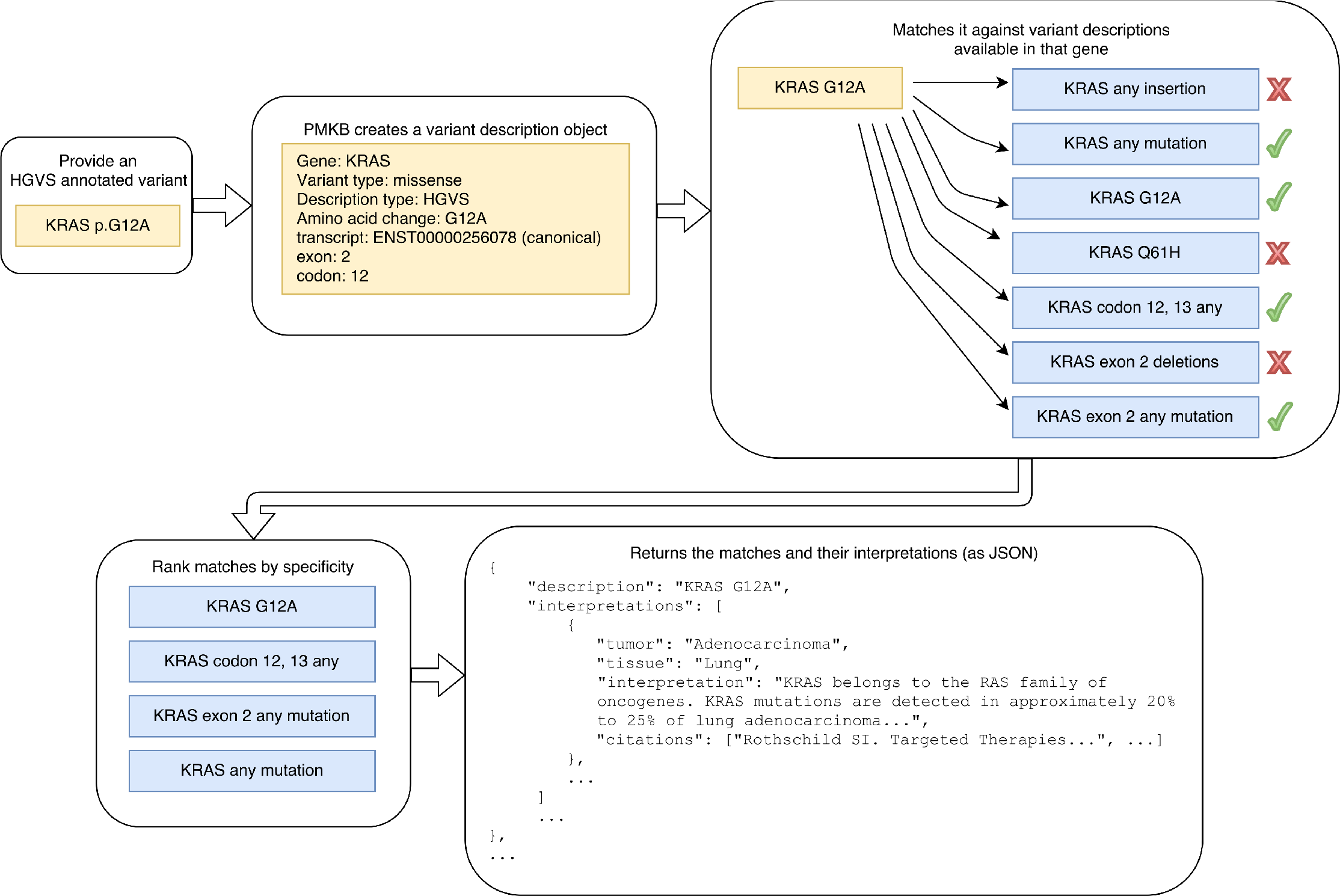
An illustration of information retuned by the PMKB API when querying a variant.

### Tumor type and tissue type objects

Tumor types correspond to possible diagnoses, such as melanoma or adenocarcinoma. Tissue types correspond to the primary site of a diagnosis. We have adopted a standard terminology for both tumor types and tissue types that was assembled by a team of highly experienced molecular pathologists. As part of the clinical genomics reporting process, patients' diagnoses need to be assigned according to this standardized terminology (such terminology may co-exist with a free-text diagnosis). These lists of tumor types (https://pmkb.weill.cornell.edu/tumors) and tissues (https://pmkb.weill.cornell.edu/tissues) are expanded and edited as needed to accurately categorize new and evolving diagnoses.

## RESULTS

### User Interface

The PMKB currently consists of a multi-user interfaces for entering, editing, browsing and querying variants. Entering variant descriptions into PMKB is done via an HTML form. For convenience, the user may enter a COSMIC ID and auto-fill the form with an HGVS description from a locally-stored version of the COSMIC database (10). Otherwise, users may choose a gene from a drop-down list or type text in a search box to search for any gene symbol accepted by the Human Genome Organization (HUGO) (11). Once a gene is chosen, the user will choose a variant type, which determines what other fields are required. For small and localized mutations, the user can choose a description type: HGVS notation, codon, exon, or “anywhere in gene”, bringing up a field with the HGVS notation, codon range, or exon range respectively. Additionally, the user may flag a variant description as germline using a separate checkbox. If “CNV” is selected as the variant type, the user will have to specify either gain or loss, and can select a checkbox if they wish to use a chromosome-based location instead of a gene-based location. Chromosome-based locations use extra fields for the chromosome and cytoband. Choosing “rearrangement/fusion” for the variant type will bring another field for partner gene that also allows choosing from any HUGO symbol. Finally, the user must provide a versioning comment that will be preserved in the change log for that entry. Once a variant is submitted, PMKB automatically adds region information using Ensembl’s API that is useful to put variants in the proper standardized context.

The user interface for entering interpretations also uses an HTML form (**Figure 3a**). This form allows the user to first select a gene from any gene in PMKB that has at least one variant description. The user will then see a list of variant descriptions for that gene, and can check off one or as many as are relevant. This is a very powerful feature that greatly facilitates applying a single interpretation to many variants, including new variants that are encountered on a rolling basis which warrant the same interpretive comment. In addition, this feature also allows the user to easily edit/modify an interpretive comment already present in the knowledgebase and apply that newly edited comment to one or more specific variants among the many variants present in the knowledgebase for that gene. Tumor types and tumor sites are also available in drop-down lists, and users may add as many entries as they see fit. Radio buttons are used to select a tier, and the actual text of the interpretation goes into a specific text field. Citations are entered using PubMed IDs, one per line in a text box. After the interpretation is submitted, the PubMed API will be used to turn these IDs into citation strings, allowing for greater consistency and convenience in citation formatting (**Figure 3b**). All interpretations must be supported by at least one literature citation. Finally, the user should enter a versioning comment that will be preserved in the entry’s change log. Interpretation pages provide links to associated variants, tumor types, tissue types, and PubMed entries, including links to external online resources such as Ensembl, COSMIC, PubMed, etc.

**Figure 3:**
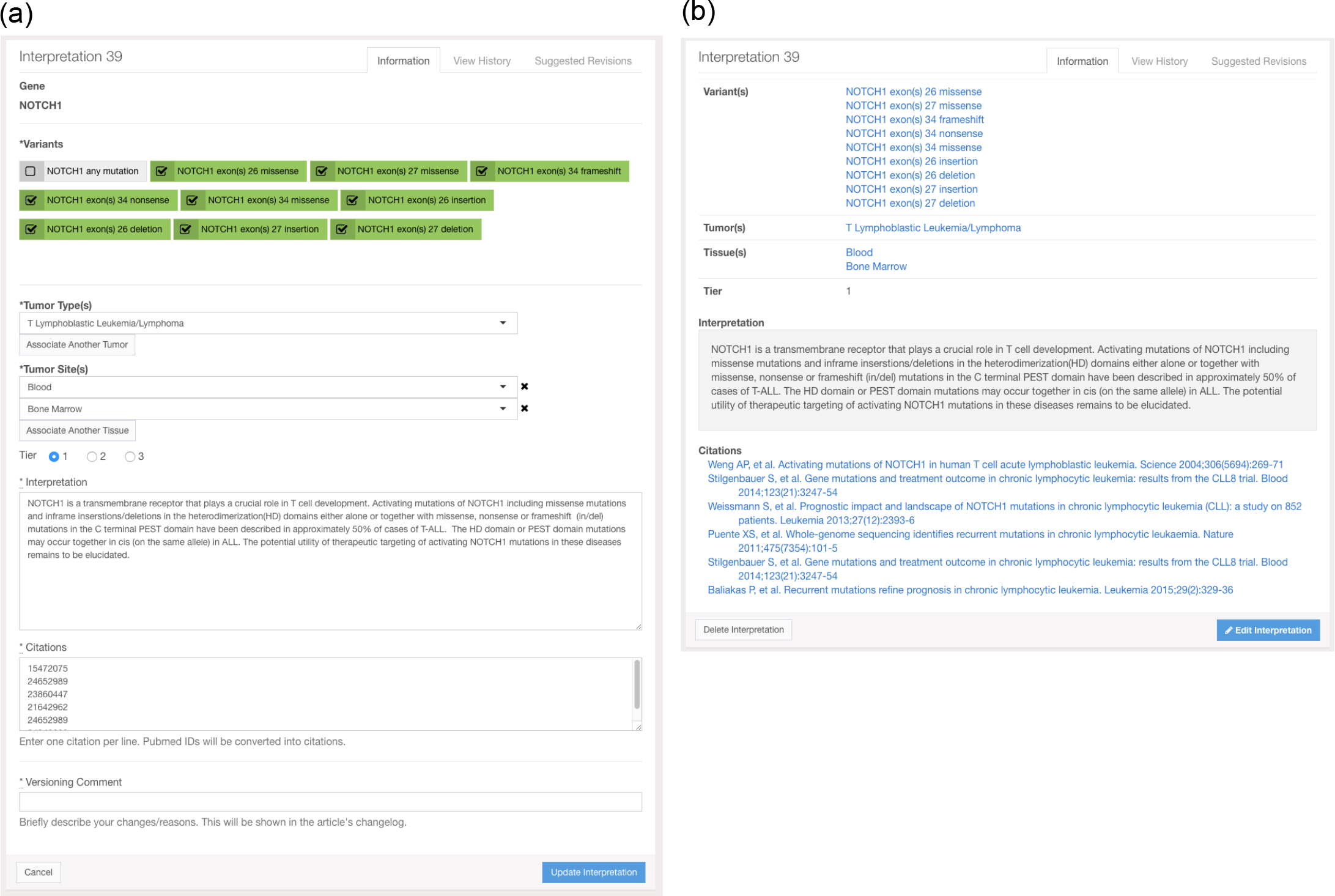
Screenshots of (a) the interface for entering an interpretation in PMKB and (b) summary view of an interpretation after entry/edit. This also demonstrates how PubMed IDs in the citations entry were resolved to complete citations in the display.

A search engine with auto-complete function can help query the PMKB for specific genes and variants. PMKB’s repository of information has been growing steadily thanks to continuous curation efforts by our molecular pathology team (**Figure 4**). At the time of writing, PMKB contains 457 variant descriptions with 281 interpretations (**Figure 4**). Genes including EGFR, BRAF, KRAS, and KIT are associated with the largest numbers of interpretable variants (**Figure 5a and 5b**). Adenocarcinomas are by far the largest tumor type, followed by acute myeloid leukemia and myelodysplastic syndromes (**Figure 5c**). The usage pattern and content are a result of the tumor types and tissue types that are currently being tested by the different platforms at our institution, which are expected to expand over time.

**Figure 4:**
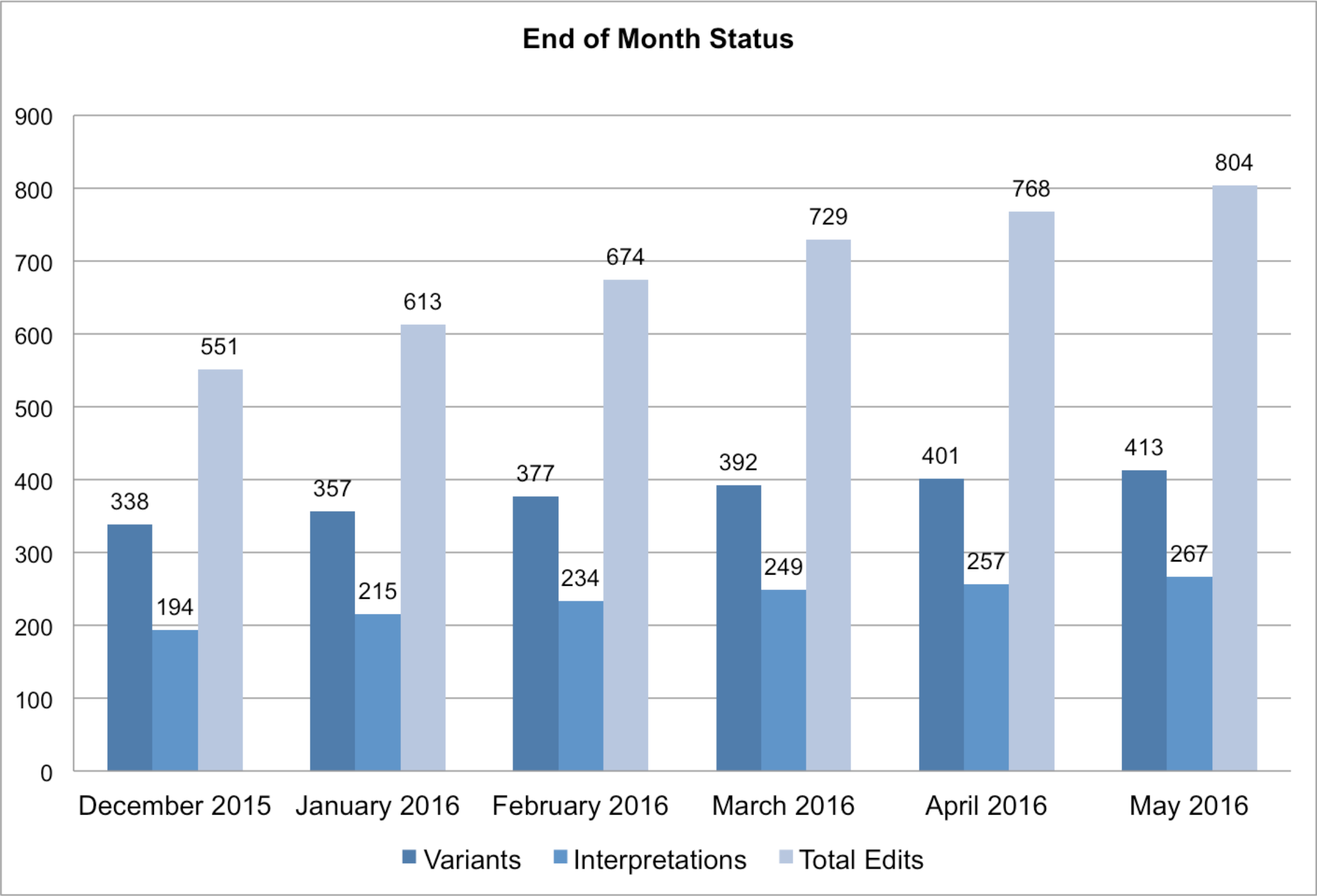
Growth of the knowledgebase over time. Entries created prior to December 2015 represent work that IPM pathologists had saved in Excel spreadsheets. Currently, a small team at the IPM makes contributions to PMKB, and expansion in PMKB’s userbase could result in a much higher rate of growth.

**Figure 5:**
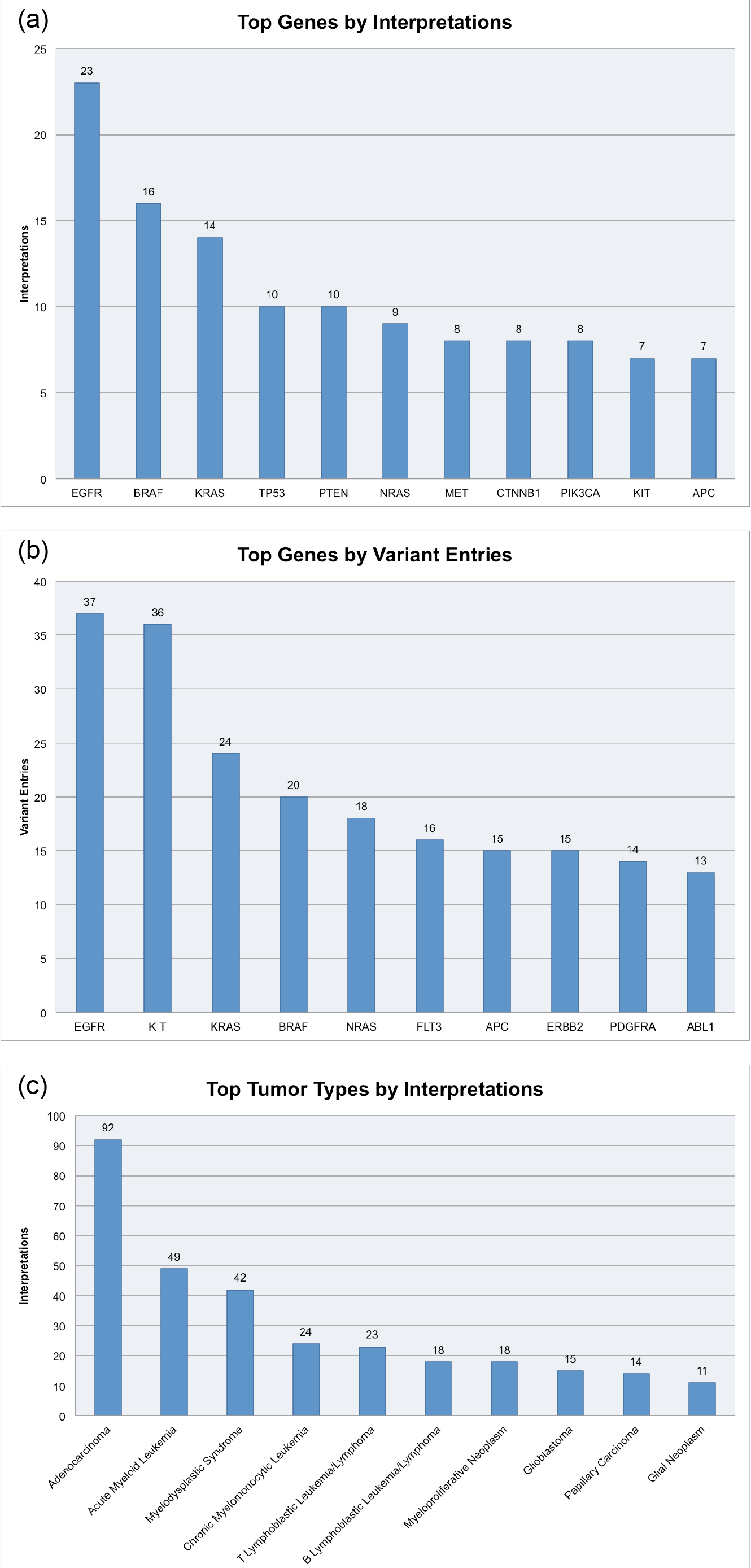
(a) The top ten genes in PMKB when ranked by number of variant descriptions in the database, (b) The top ten genes in PMKB when ranked by number of interpretations in the database. The prevalence of certain genes in PMKB corresponds naturally to genes that have well-studied variants with possible diagnostic use. (c) The top ten solid tumor types in PMKB when ranked by number of interpretations.

### User roles

Within the PMKB application, users have three different levels of privilege - a high-level “approver” who can review and approve others’ entries, standard users who can submit edits, and guests who cannot make changes. The first role is reserved for the PMKB’s molecular pathologists, who are able to enter any changes to variant descriptions or interpretations directly for publication on the site. The second role is intended for general users, such as clinical fellows, medical students, members of the computational team, etc., who may create and edit variant descriptions and interpretations, but their new entries and modifications must be approved by an approver pathologist before being published on the site. A pathologist may also choose to edit another user’s entry before approval. Since interpretations are used for clinical reporting, signing-out molecular pathologists can be guaranteed that all changes have been approved by a board-certified pathologist. All changes are also tracked in an audit log for each entry. Administrators and users in the first-tier role may restore any variant description or interpretation to a previous state based on the audit log. Auditing is an important part of clinical workflows, and PMKB is able to provide this capability in a robust capacity.

For potential collaborations outside of WCM, Security Assertion Markup Language (SAML)-based authentication allows a wider variety of users to access and make edits to PMKB. SAML is a format that allows for standardized exchange of authentication data, providing the capability to use institution-based logins within external web applications. WCM is part of the SAML-based InCommon federation, and the registration of PMKB as an InCommon website opens up PMKB to collaborations with pathologists and researchers at other institutions. As curating interpretations is a common activity related to genomic testing reports, this could potentially reduce repetition of that work between institutions.

### An example of API integration

To illustrate integration with PMKB’s API and the use of PMKB to retrieve interpretations, we describe a tool called the AmpliSeq Results Converter that was developed in the IPM specifically to aid processing of variants in VCF format for the AmpliSeq assay (Ion AmpliSeq™ Cancer Hotspot Panel v2, Life Technologies). The AmpliSeq Results Converter converts annotated VCFs into more human-friendly spreadsheets and text reports to facilitate reporting in the electronic health record. The AmpliSeq Results Converter is a Python script that queries the PMKB based on a sample’s tumor and primary site, and variants’ HGVS annotations within the VCF. Queries occur via the PMKB’s REST API through HTTP requests. PMKB returns the most relevant interpretation for each request and the Ampliseq Results Converter generates a report suitable for upload into the laboratory information systems (LIS) Cerner Millenium and Helix. The AmpliSeq Results Converter has greatly facilitated the laboratory’s workflow by reducing the need to manually copy interpretations from an Excel spreadsheet into the diagnostic report. The use of this pipeline allows the appropriate interpretations to be pulled automatically from the PMKB into Millenium Helix, our clinical reporting system. All final reports are reviewed by molecular pathologists prior to case sign-out to ensure that the proper associations between tumor type, tissue type, gene variant and interpretive comment are made. The lab uses PMKB as its primary tool for entering and retrieving interpretation information, starting in late December 2015. This use of PMKB’s API has significantly facilitated the process of retrieving interpretations for variants, especially when the interpretations were generalized for gene regions (e.g. EGFR exon 19) rather than specific point mutations (e.g. EGFR p.L858R). Currently, PMKB’s API for interpretation retrieval has been used for several hundred cases and the PMKB is used routinely as part of the clinical workflow.

## CONCLUSION

We have herein described the Precision Medicine Knowledge Base, an interactive online application for collaborative editing, maintenance and sharing of structured clinical-grade cancer mutations interpretations. All interpretations are made available free-of-charge to the community and can be accessed either via the Web interface (https://pmkb.weill.cornell.edu) or programmatically via the existing API. Within our institution, the PMKB has already proven to be an enormously useful tool for storing and retrieving interpretation information amenable both to use by pathologists and reporting pipelines. It has led to significant improvements in the laboratory workflow for the Ampliseq 50-gene panel assay, saving considerable time and effort, and is currently being used to report WES results (3).

Since many institutions face similar problems with reporting, it is our hope that the success of PMKB at WCM/NYPH can be replicated and expanded with potential collaborators at other institutions. To support both larger assays in-house and potential traffic from collaborators, PMKB’s infrastructure, SAML authentication capabilities, and API can readily be leveraged, while continuing to serve results reliably at acceptable speeds. The speed of computation becomes highly relevant with larger panels of genes that may return calls for hundreds of different variants, an issue that can be addressed with further parallel processing and database query optimization within PMKB’s API.

## DISCUSSION

In addition to improving PMKB’s technical performance, there is the challenge of increasing the output of the interpretations themselves. Quality and quantity of interpretations is what will make PMKB attractive to potential collaborators, and will set a standard for future contributions to the knowledgebase. Since interpretations must be written by qualified individuals, the PMKB software tries to ease the burden of storing and organizing interpretations, allowing pathologists to focus on writing interpretive comments and signing out cases. Along with writing new interpretations comes the labor of keeping interpretations and tier information up-to-date with current information on published literature, clinical trials, and FDA drug approval. Bringing in clinicians and physicians, currently only on the receiving end of the reporting process, collaborators, and outside contributors could significantly increase the output, quality, and maintainability of interpretations, but this activity must be managed carefully within PMKB, to ensure that all edits are up to par. It is important to make sure that PMKB has tools for managing users and user-produced content to prepare for a growing user base.

PMKB will need to continue to adapt based on feedback from users, in order to make sure a variety of needs around reporting are met. Some examples are tumor type specific pertinent negatives (already supported in PMKB but not yet systematically annotated) and ongoing improvements to interpretation content. Another future feature includes adding a “talk page” style interface for users to communicate with one another about potential edits and entries, which will become important to track discussion in an expanded user base, especially when an interpretation needs to be brought up-to-date with new literature. Extensions of the API will provide more functions for retrieving interpretations and meet some of the ongoing issues described above. Ideally, this Knowledgebase will serve as an open tool amenable to crowd-sourcing of content over time by experts in specific subspecialties.

## AKNOWLEDGEMENTS, COMPETING INTERESTS, FUNDING

The authors would like to thank Elemento lab and Institute for Precision Medicine members for their feedback and discussions. No competing interests are reported. This work was supported by the CAREER grant from National Science Foundation (DB1054964), NIH grant R01CA194547, the Starr Cancer Foundation, as well as by startup funds from the Institute for Computational Biomedicine.

## Supplemental figures

**Figure S1:**
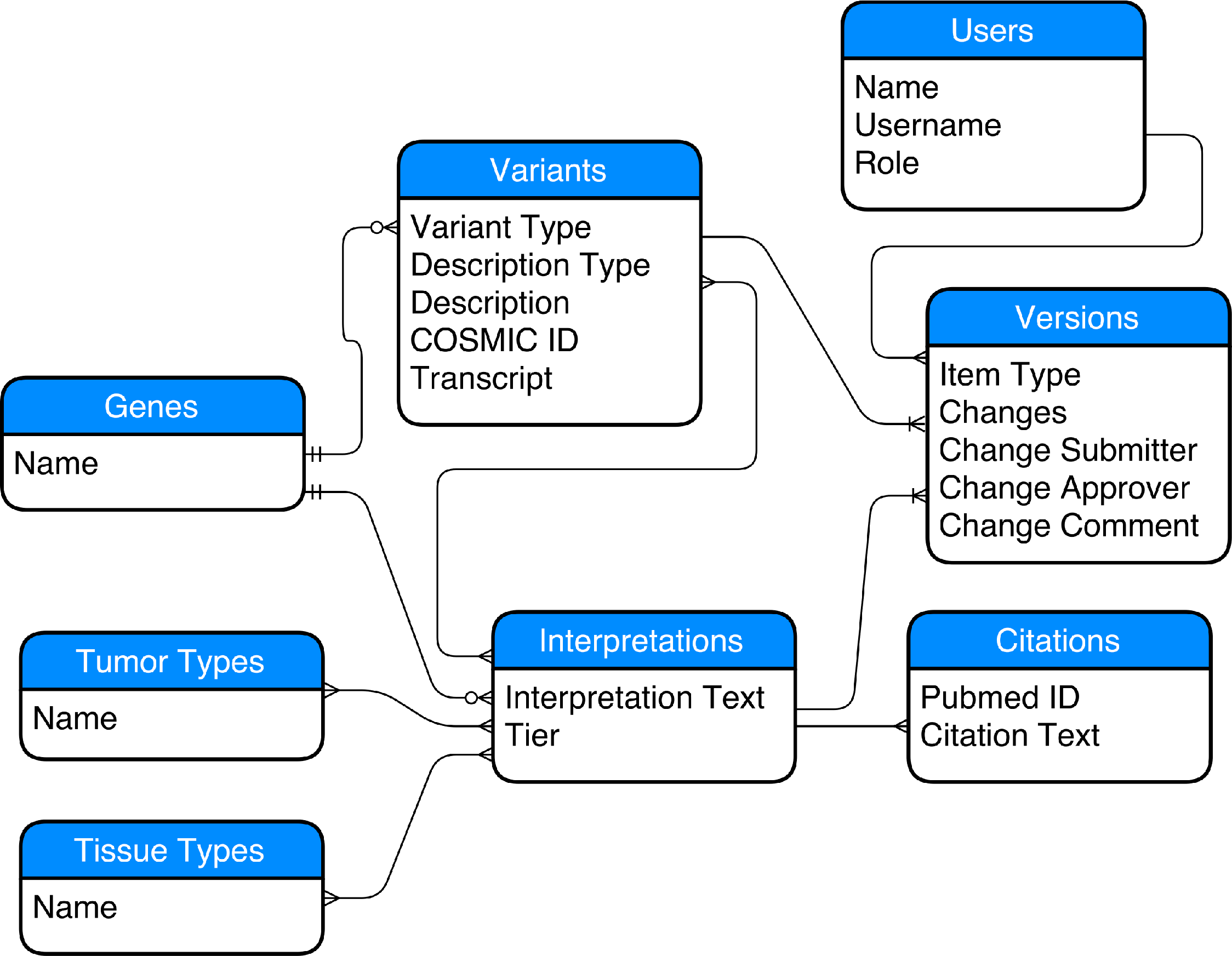
A simplified entity-relationship (ER) diagram of PMKB’s database (not showing all table columns). PMKB uses a relational database, a design that reinforces unique data that can be reused through multiple pre-defined associations with other data objects. Separate tables for tumor types, tissue types, variants, interpretations, and citations allows for more flexibility with associations.

